## Supplementary figures and tables for "Caught in the Act: targeted mutagenesis of the herpesvirus fusogen central helix captures transition states"

**For**

Zhou et al.

| Genus | Virus | Start | Sequence | End | Host species | NCBI accession |
|---|---|---|---|---|---|---|
| Varicellovirus | VZV | 511- | SVEFAMLQFTYDHIQ**EHV**NEMLARISSSWCQLQNRERALWSGLFPI | -556 | Human | AGC94521 |
| | CeHV-9 | 501- | SVEFAMLQFTYDHIQ**EHV**NEMLARISTSWCQLQNKERALWSSLFPV | -546 | African green monkey | AAA58703 |
| | PRV | 528- | SAEFARLQFTYDHIQ**AHV**NDMLSRIAAAWCELQNKDRTLWGEMSRL | -573 | Pig | ALB06102 |
| | EHV-1 | 574- | SIEFAMLQFAYDHIQ**SHV**NEMLSRIATAWCTLQNKERTLWNEMVKI | -619 | Horse | BAF43295 |
| | EHV-3 | 590- | SVEFAMLQFAYDHIQ**AHV**NEMLSRIATAWCTLQNKERVLWNEMVKI | -635 | Horse | YP_009054936 |
| | EHV-4 | 569- | SIEFAMLQFAYDHIQ**SHV**NEMLSRIATAWCTLQNKERTLWNEMVKV | -614 | Horse | QYL35303 |
| | EHV-8 | 573- | SIEFAMLQFAYDHIQ**SHV**NEMLSRIATAWCTLQNKERTLWNEMVKI | -618 | Horse | AUS94926 |
| | EHV-9 | 475- | SIEFAMLQFAYDHIQ**SHV**NEMLSRIATAWCTLQNKERTLWNEMVKI | -520 | Horse | YP_002333514 |
| | FeHV-1 | 538- | SVEFAMLQFAYDYIQ**AHV**NEMLSRIATAWCTLQNREHVLWTETLKL | -583 | Cat | ALJ85888 |
| | CaHV-1 | 472- | SVHFAMLQFAYDHIQ**SHV**NEMLSRIATAWCNLQNKERTLWNEVMKL | -517 | Dog | AEK27122 |
| | BoHV-1 | 535- | SAEFAALQFTYDHIQ**DHV**NTMFSRLATSWCLLQNKERALWAEAAKL | -580 | Cattle | QBI59636 |
| | BoHV-5 | 541- | SAEFAALQFTYDHIQ**DHV**NTMFSRLATSWCLLQNKERALWAEAAKL | -586 | Cattle | ARM56009 |
| | BuHV-1 | 545- | SAEFAALQFTYDHIQ**DHV**NTMFSRLATSWCLLQNKERALWAEAAKL | -590 | Buffalo | APO15888 |
| | CpHV-1 | 519- | SAEFAALQFTYDHIQ**DHV**NAMFSRLATSWCLLQNKERALWAEAAKL | -564 | Goat | YP_009664694 |
| | CvHV-1 | 530- | SAEFAALQFTYDHIQ**DHV**NTMFSRLATSWCLLQNKERALWAEAAKL | -575 | Deer | YP_009664695 |
| | CvHV-2 | 532- | SAEFAALQFTYDHIQ**DHV**NAMFSRLATSWCQLQNKERTLWAEAAKL | -577 | Deer | AVT50752 |
| | CvHV-3 | 534- | SAEFAALQFTYDHIQ**DHV**NTMFSRLATSWCLLQNKERALWAEAAKL | -579 | Deer | YP_010087575 |
| | MoHV-1 | 502- | GAEFAALQFTYNHIQ**SHV**NEMFARIATAWCALQNKELALWKETVKL | -547 | Beluga whale | YP_010084963 |
| | PhHV-1 | 478- | SIHFAMLQFAYDHIQ**SHV**NEMLSRIATAWCNLQNKERTLWNEVMKL | -523 | Seal | CAA92272 |
| Simplexvirus | HSV-1 | 500- | SIEFARLQFTYNHIQ**RHV**NDMLGRVAIAWCELQNHELTLWNEARKL | -545 | Human | AFH41177 |
| | HSV-2 | 497- | SIEFARLQFTYNHIQ**RHV**NDMLGRIAVAWCELQNHELTLWNEARKL | -542 | Human | CAB06752 |
| | CeHV-2 | 485- | SVEFARLQFTYDHIQ**RHV**NDMLGRIATAWCELQNRELTLWNEARRL | -530 | African green monkey | CAA40256 |
| | AtHV-1 | 523- | TVEFARLQFTYDHIQ**KHV**NEMLGRIAAAWCQLQNQELVLWNEARKL | -568 | Spider monkey | YP_009361910 |
| | McHV-1 | 487- | SVEFARLQFTYDHIQ**RHV**NDMLGRIAIAWCELQNHELTLWNEARKL | -532 | Macaque monkey | NP_851887 |
| | PnHV-3 | 499- | SIEFARLQFTYNHIQ**RHV**NDMLGRIAVAWCELQNHELTLWNEARKL | -544 | Chimpanzee | YP_009011014 |
| | SaHV-1 | 509- | SVEFARLQFTYDHIQ**KHV**NEMLGRIAAAWCQLQNQELVLWNEARKL | -554 | Squirrel monkey | AAM22795 |
| | PaHV-2 | 488- | SIEFARLQFTYDHIQ**RHV**NDMLGRIAIAWCELQNRELTLWNEARRL | -533 | Baboon | BAE79592 |
| | BoHV-2 | 518- | SVEFARLQFTYNHIQ**KHV**NEMFGRMAVSWCELQNQELTLWNEAKKI | -563 | Cattle | YP_009664618 |
| | LeHV-4 | 482- | SVEFARLQFTYDHIQ**RHV**NEMLGRLAVSWCELQNQELTLWNEAQKL | -527 | Rabbit | YP_009230158 |
| | MaHV-1 | 483- | SIEFARLQFTYDHIQ**QHV**NEMLGRIAIAWCELQNRELTLWNEARKL | -528 | Parma wallaby | YP_009227260 |
| | MaHV-2 | 486- | SIEFARLQFTYDHIQ**QHV**NEMLGRIAIAWCELQNRELTLWNEARKL | -531 | Quokka | YP_009664622 |
| | PtHV-1 | 340- | SIEFARLQFTYDHIQ**KHV**NDMLGRIAIAWCQLQNQELVLWNEARKL | -385 | Fruit bat | YP_009042089 |
| Mardivirus | AnHV-1 | 586- | SAQFAMLQYTYDHIQ**AHV**NDMLSRIAVSWCELQNKESVLWAEMRKV | -631 | Duck | AET71141 |
| | CoHV-1 | 579- | SAQYAMLQFTYDHIQ**GHV**NDMFSRIAVAWCELQNKERVLWSEALKI | -624 | Pigeon | YP_009352935 |
| | MeHV-1 | 463- | SVQFAMLQFLYDHIQ**AHI**NEMFSRIATAWCELQNKELVLWREAIKI | -508 | Turkey | NP_073321 |
| | GaHV-2 | 457- | SVQFAMLQFLYDHIQ**THI**NDMFSRIATAWCELQNRELVLWHEGIKI | -502 | Chicken | AAM97702 |
| | GaHV-3 | 458- | SVQFAMLQFLYDHIQ**THI**NDMFSRIATAWCELQNKELALWQEGMKI | -503 | Chicken | NP_066859 |
| | SpAHV-1 | 487- | TVHFAMLQFTYDHIQ**RHV**NEMLGRIAKAWCELQNRESVLWREMRKM | -532 | Penguin | YP_009342376 |
| Iltovirus | GaHV-1 | 452- | SATFAMLQFAYDKIQ**AHV**NELIGNLLEAWCELQNRQLIVWHEMKKL | -497 | Chicken | CAA39573 |
| | PsHV-1 | 470- | SATFAMLQFTYDKIQ**KHV**NMLIGNLLEAWCEMQNRQLVVWHEIKKL | -515 | Parrot | AAQ73706 |
| Scutavirus | ChHV-5 | 431- | DATFTQVQFTYDVLR**QHI**NSVFSRLVNAWCELQNRDVTVWELLLNI | -476 | Sea turtle | QHH25918 |
| | TeHV-3 | 427- | TAAFLQIQYTYDKLQ**AHN**NAMFSRIVYAWCELQNRDITMWEQLNKI | -472 | Tortoise | YP_009176910 |
|  |  |  | ` : :*: *: :: * * ::..: :** :**:: :* :` |  |  |  |

**Figure S1. Sequence alignment of the amino acids equivalent to VZV gB DIII central helix from the five genera of the subfamily alphaherpesvirinae.** Virus host species and the glycoprotein gB NCBI accession are listed and the residues corresponding to $^{526}$EHV$^{528}$ are colored in blue. Conservation among residues is presented based on Clustal Omega: "." indicates weak similarity; ":" indicates strong similarity; "*" indicates full conservation. Histidine corresponding to H527 in VZV is highlighted in pink box. VZV: Varicella-zoster virus; CeHV-9: Cercopithecine alphaherpesvirus 9; PRV: pseudorabies virus; EHV-1: Equine alphaherpesvirus 1; EHV-3: Equine alphaherpesvirus 3; EHV-4: Equine alphaherpesvirus 4; EHV-8: Equine alphaherpesvirus 8; EHV-9: Equine alphaherpesvirus 9; FeHV-1: Felid alphaherpesvirus 1; CaHV-1: Canid alphaherpesvirus 1; BoHV-1: Bovine alphaherpesvirus 1; BoHV-5: Bovine alphaherpesvirus 5; BuHV-1: Bubaline alphaherpesvirus 1; CpHV-1: Caprine alphaherpesvirus 1; CvHV-1: Cervid alphaherpesvirus 1; CvHV-2: Cervid alphaherpesvirus 2; CvHV-3: Cervid alphaherpesvirus 3; MoHV-1: Monodontid alphaherpesvirus 1; PhHV-1: Phocid alphaherpesvirus 1; HSV-1: Herpes simplex virus 1; HSV-2: Herpes simplex virus 2; CeHV-2: Cercopithecine alphaherpesvirus 2; AtHV-1: Ateline alphaherpesvirus 1; McHV-1: Macacine alphaherpesvirus 1; PnHV-3: Panine alphaherpesvirus 3; SaHV-1: Saimiriine alphaherpesvirus 1; PaHV-2: Papiine alphaherpesvirus 2; BoHV-2: Bovine alphaherpesvirus 2; LeHV-4: Leporid alphaherpesvirus 4; MaHV-1: Macropodid alphaherpesvirus 1; MaHV-2: Macropodid alphaherpesvirus 2; PtHV-1: Pteropodid alphaherpesvirus 1; AnHV-1: Anatid alphaherpesvirus 1; CoHV-1: Columbid alphaherpesvirus 1; MeHV-1: Meleagrid alphaherpesvirus 1; GaHV-2: Gallid alphaherpesvirus 2; GaHV-3: Gallid alphaherpesvirus 3; SpAHV-1: Spheniscid alphaherpevirus 1; GaHV-1: Gallid alphaherpesvirus 1; PsHV-1: Psittacid alphaherpesvirus 1; ChHV-5: Chelonid alphaherpesvirus 5; TeHV-3: Testudinid alphaherpesvirus 3.

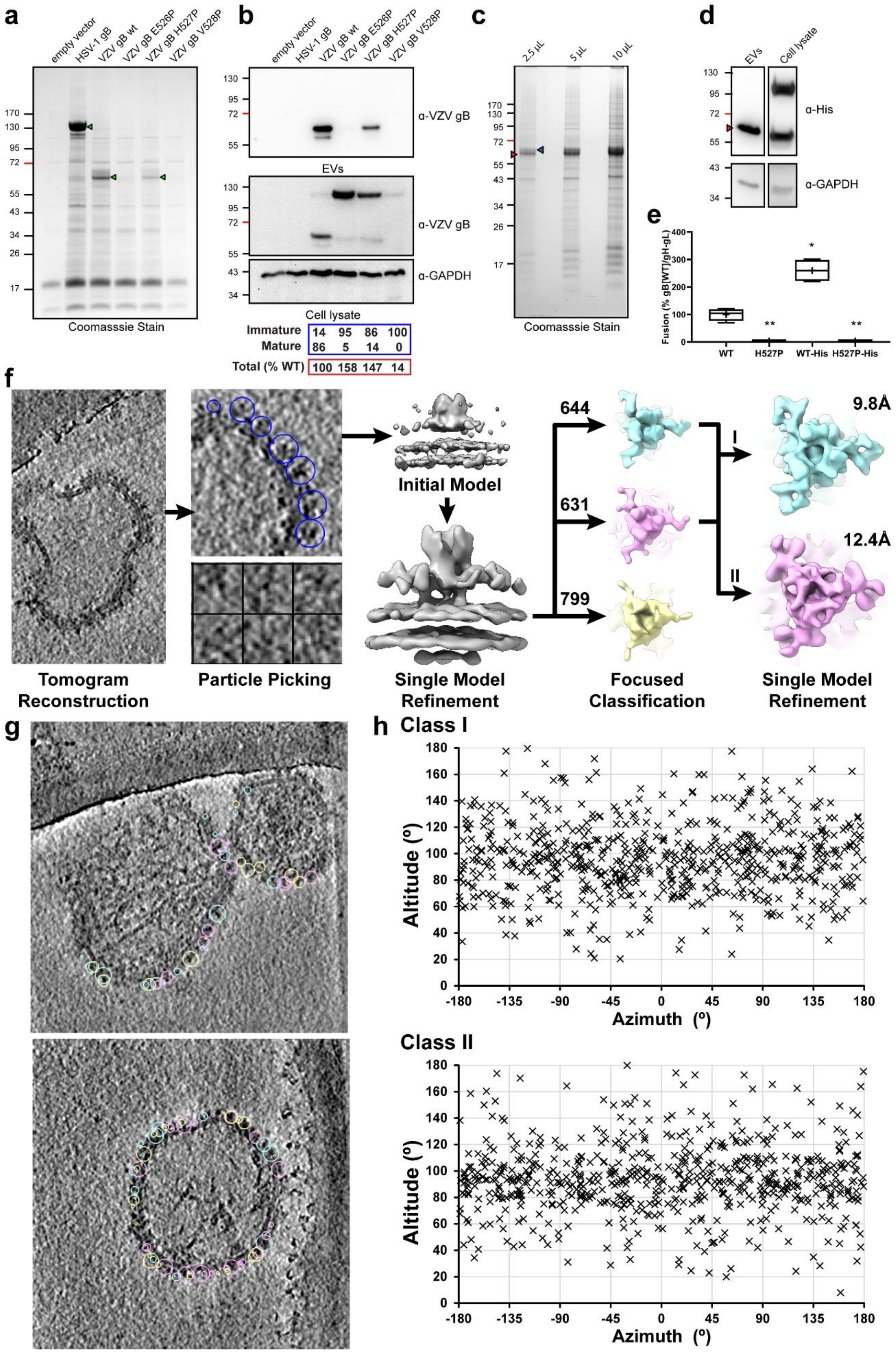
a
empty vector
HSV-1 gB
VZV gB wt
VZV gB E526P
VZV gB H527P
VZV gB V528P
Coomasssie Stain
b
EVs
α-VZV gB
α-GAPDH
Cell lysate
Immature 14 95 86 100
Mature 86 5 14 0
Total (% WT) 100 158 147 14
c
2.5 μL
5 μL
10 μL
Coomasssie Stain
d
EVs
Cell lysate
α-His
α-GAPDH
e
Fusion (% gB[WT]/gH-gL)
WT
H527P
WT-His
H527P-His
f
Tomogram Reconstruction
Particle Picking
Initial Model
Single Model Refinement
644
631
799
Focused Classification
I
II
9.8Å
12.4Å
Single Model Refinement
g
h
Class I
Class II
Altitude (º)
Azimuth (º)

**Figure S2. Characterization of EVs expressing gB proline substitutions to H527 residue in DIII. a** Coomassie blue stain of EVs purified from BHK-21 cells transfected with vectors expressing VZV WT gB, gB $^{526}$EHV$^{528}$ proline mutants, HSV-1 gB or empty vector. Green arrow heads highlight the uncleaved form of HSV-1 gB and the cleaved form of VZV gB. **b** Western blot of EVs from (**a**) detected with rabbit anti-VZV gB 746-867 and whole cell lysates from transiently transfected BHK-21 cells; GAPDH loading control. The relative molecular abundance of the immature full-length, and the mature cleaved fragment of gB in the transfected cells were shown as a percentage of the sum abundance of the two gB forms (blue box); the combined band density of the immature full-length and the cleaved fragment of gB in cell lysates was normalized to that of GAPDH and shown as a percentage of WT gB (red box). **c** Coomassie blue stain of EVs purified from BHK-21cells transfected with vectors expressing the C-terminally His-tagged gB[H527P] (H527P-His). The SDS-PAGE gel was loaded with increasing quantities of EVs. N-terminal and C-terminal cleavage products of VZV gB[H527P] are indicated by blue and red arrowheads respectively. **d** Western blot of EVs from (**c**) and cell lysates detected for gB (anti-His tag); GAPDH loading control; C-terminal cleavage product of VZV gB[H527P] is indicated by red arrowhead. Mass markers (kDa) are shown on the left (**a-d**). **e** Box and whisker plots for cell-cell fusion measured by the stable reporter fusion assay (SRFA) using CHO-DSP1 cells transfected with vectors expressing untagged (WT) or His-tagged WT gB (WT-His) or the corresponding H527P mutants (H527P and H527P-His) with gH-gL, and mixed with MeWo-DSP2 cells for 48hrs. Fusion efficiency was normalized to that mediated by gB[WT]/gH-gL (%). Box represents 25-75 percentile, whiskers extend to 10-90 percentile, the median is the horizontal band and the mean is the **+**. Data from two independent experiments is shown. The statistical difference compared to WT gB was analyzed by one-way ANOVA (*, $p < 0.1$; **, $p < 0.01$). **f** Classification scheme of gB[H527P] particles in EVs membranes identified in cryo-ET micrographs, highlighted blue circles, initially yielded three classes (Cyan: Class I; Purple: Class II; Yellow: Class III). Further analysis dismissed Class III due to low resolution in the central region, finally yielding the two classes producing the Class I (9.8Å) and Class II (12.4Å) cryo-EM maps. Particle numbers for each class are shown for the Focused Classification. **g** Distribution of Class I to III particles on the surface of EVs. The circles represent manual particle picking for each class, color corresponding to each class shown in (**f**). **h** Angular distribution of the particles used to generate the Class I and Class II Cryo-EM maps.

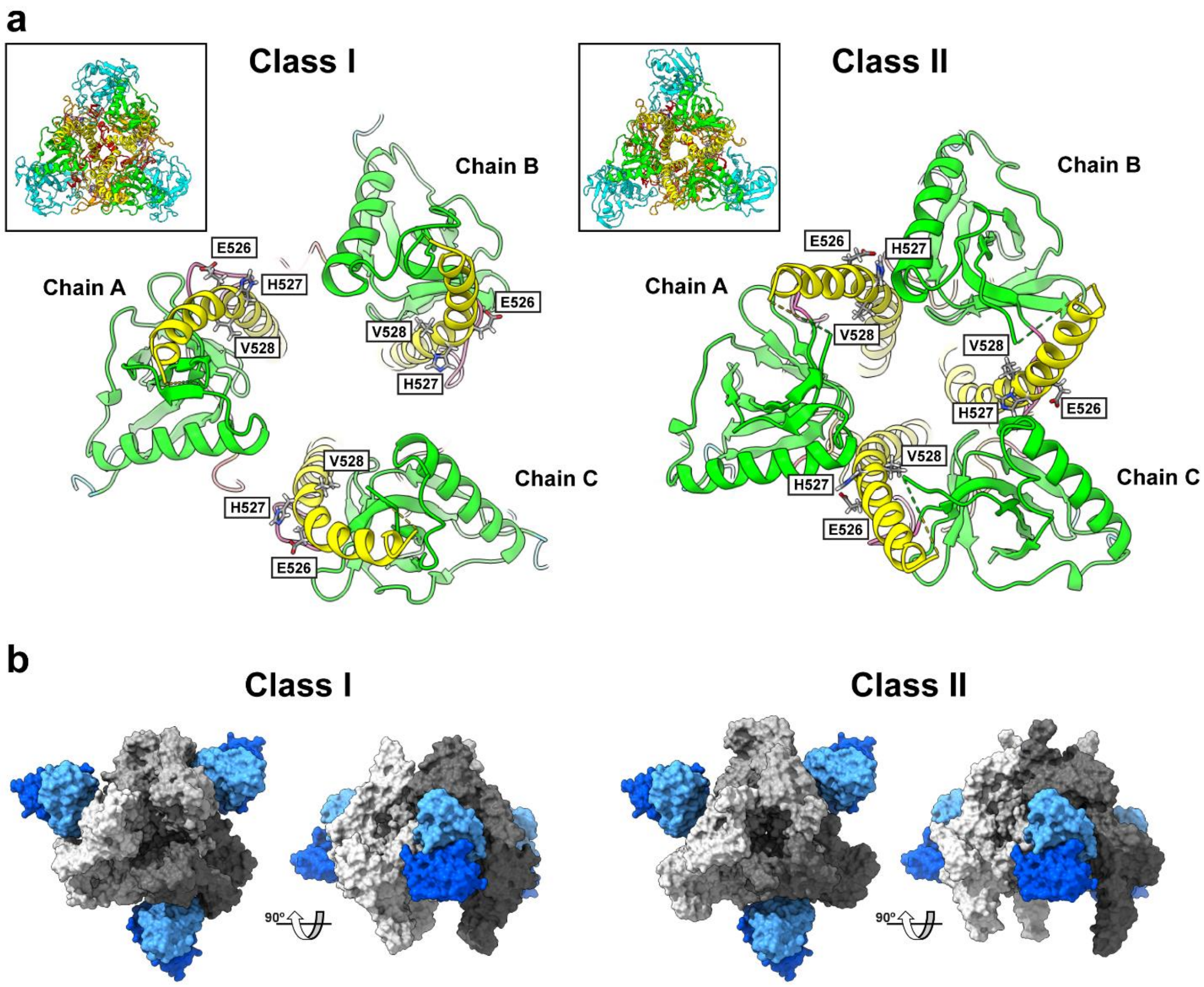

**Figure S3. The location of residues $^{526}$EHV$^{528}$ and predicted binding of mAb 93k for Class I and Class II models of gB[H527P]. a** Ribbon diagrams highlight the locations of the $^{526}$EHV$^{528}$ residues in the gB DIII central helix. The insets show the top views of the Class I and Class II models with each domain colored; DI (cyan), DII (green), DIII (yellow), DIV (orange), DV (red) and linker regions (hot pink). The main panels show an area of the Class I and Class II models focused on DII and the upper portion of the DIII central helix for each of the chains A to C. **b** Molecular surface representation demonstrating the location of human, VZV neutralizing mAb 93k fab fragments predicted to bind to the Class I and Class II models. The three chains of VZV gB are colored white (A), grey (B), and dark grey (C), and the heavy and light chains of the mAb 93k fab fragments are colored blue and light blue respectively. Fab fragments of mAb 93k bind to VZV gB DIV [1,2].

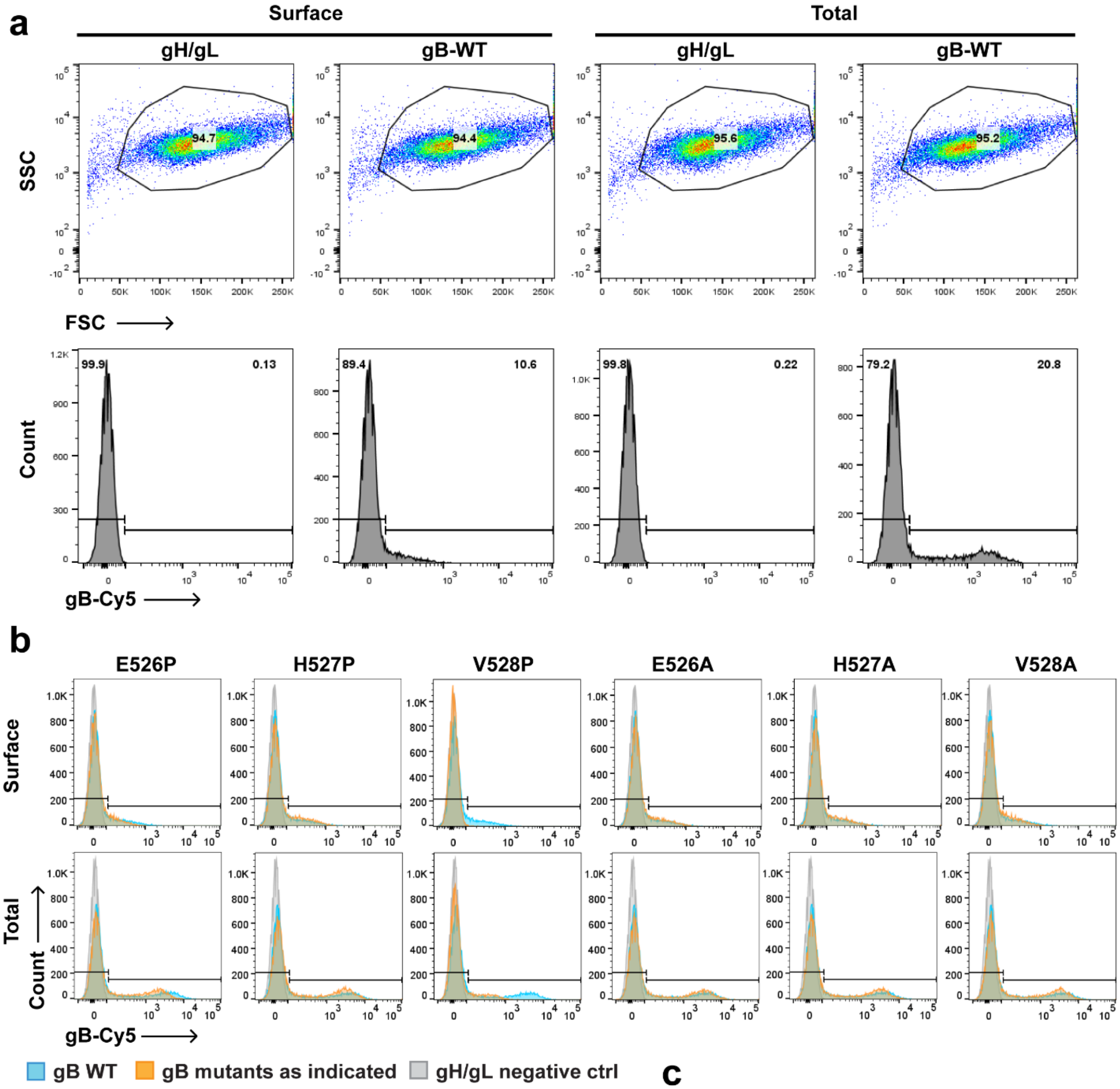

| | Frequency of postive cells (%WT) | | Median fluorescence intensity (%WT) | |
|---|---|---|---|---|
| | Surface | Total | Surface | Total |
| **E526P** | 80.5 ± 3.3 | 109.0 ± 14.9 | ND | 70.3 ± 8.9 |
| **H527P** | 119.3 ± 16.6 | 118.1 ± 14.7 | ND | 95.4 ± 3.6 |
| **V528P** | 3.9 ± 0.7 | 48.0 ± 8.2 | ND | 17.5 ± 0.6 |
| **E526A** | 119.6 ± 3.6 | 118.8 ± 3.7 | ND | 109.6 ± 0.7 |
| **H527A** | 110.8 ± 5.2 | 120.9 ± 6.3 | ND | 98.1 ± 3.6 |
| **V528A** | 104.4 ± 6.0 | 120.6 ± 2.4 | ND | 98.5 ± 6.8 |

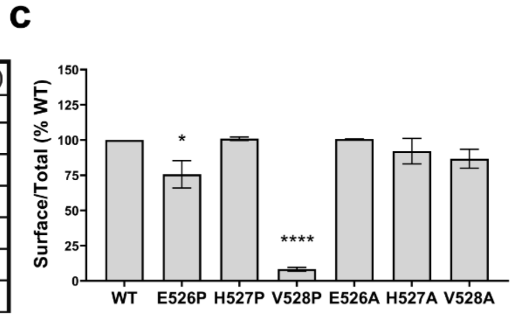

**Figure S4. The analysis of gB expression by flow cytometry. a** Gating strategy for the quantification of gB levels using mAb 93k in flow cytometry. The representative samples are for total and surface staining of CHO-DSP1 cells transfected with gH/gL or gB and detected with the anti-gB mAb Cy5-conjugated 93k. The top panels represent examples of CHO cells nonpermeabilized for surface, or permeabilized for total, gated using forward scatter (FSC) and side scatter (SSC). The lower panels represent the histograms from the gated CHO cells and show the specificity of the 93k mAb for VZV gB. **b** Surface or total expression of WT gB or gB mutants. The representative samples show the population of WT gB-expressing cells (blue) and mutant gB-expressing cells (orange) was gated against cells transfected with gH-gL (grey). The frequency of gB-expressing cells and median of fluorescence intensity (MFI) were compared to that in the WT gB and presented as percentage of WT in the table (%WT; mean ± SD). **c** The ratio of surface to total gB expression from **a** normalized to WT gB (% WT) (mean ± SEM) was calculated. Statistical differences from at least two independent experiments were evaluated by one-way ANOVA (*, $p < 0.1$; ****, $p < 0.0001$).

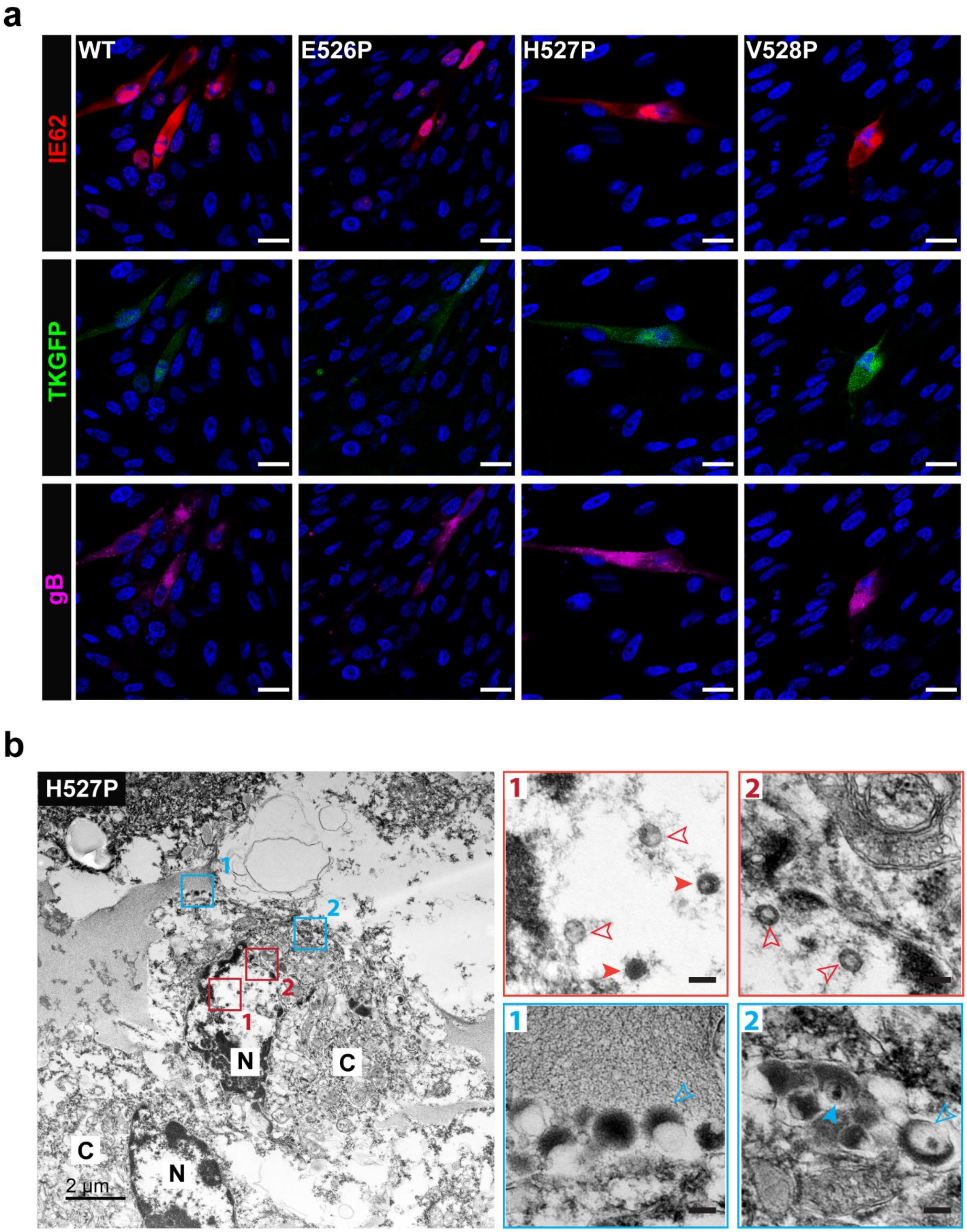
a
WT
E526P
H527P
V528P
IE62
TKGFP
gB
b
H527P
1
2
N
C
C
N
2 µm

**Figure S5. Proline substitutions to $^{526}$EHV$^{528}$ residues in DIII does not interfere with gB expression and virus assembly in BAC transfected cells. a** Confocal microscopy images of MeWo cells transfected with pOka-TKGFP (WT), pOka-TKGFP-gB[E526P] (E526P), pOka-TKGFP-gB[H527P] (H527P), or pOka-TKGFP-gB[V528P] (V528P) BACs at 72 hrs post transfection. All the BACs expressed thymidine kinase (TK) fused with GFP fluorescence protein at the C-terminus. Cells were stained for immediate-early protein (IE62; red), gB (93k; violet), and nuclei (Hoechst 33342; blue). Scale bar = 20 µm. **b** Transmission Electron Microscopy (TEM) images of MeWo cells transfected with pOka-TKGFP-gB[H527P] BAC at 48 hrs post transfection. The transfected cells were harvested at 48 hrs post transfection and enriched for GFP-expressing cells by sorting prior to TEM. Panels at the right are magnifications of areas indicated by boxes in the larger images (N: nuclei; C: cytoplasm). Boxes outlined in red show representative viral capsids, with solid red arrowheads indicating viral DNA containing C-capsid and open red arrowheads indicating empty A-, or scaffold containing B capsids. Boxes outlined in blue show representative viral particles, with solid blue arrowhead indicating complete virus particle in the process of assembly in the cytoplasm and open blue arrowheads indicating light particles on the cell surface or in the process of assembly in the cytoplasm. In the magnified images, scale bar = 0.1 µm.

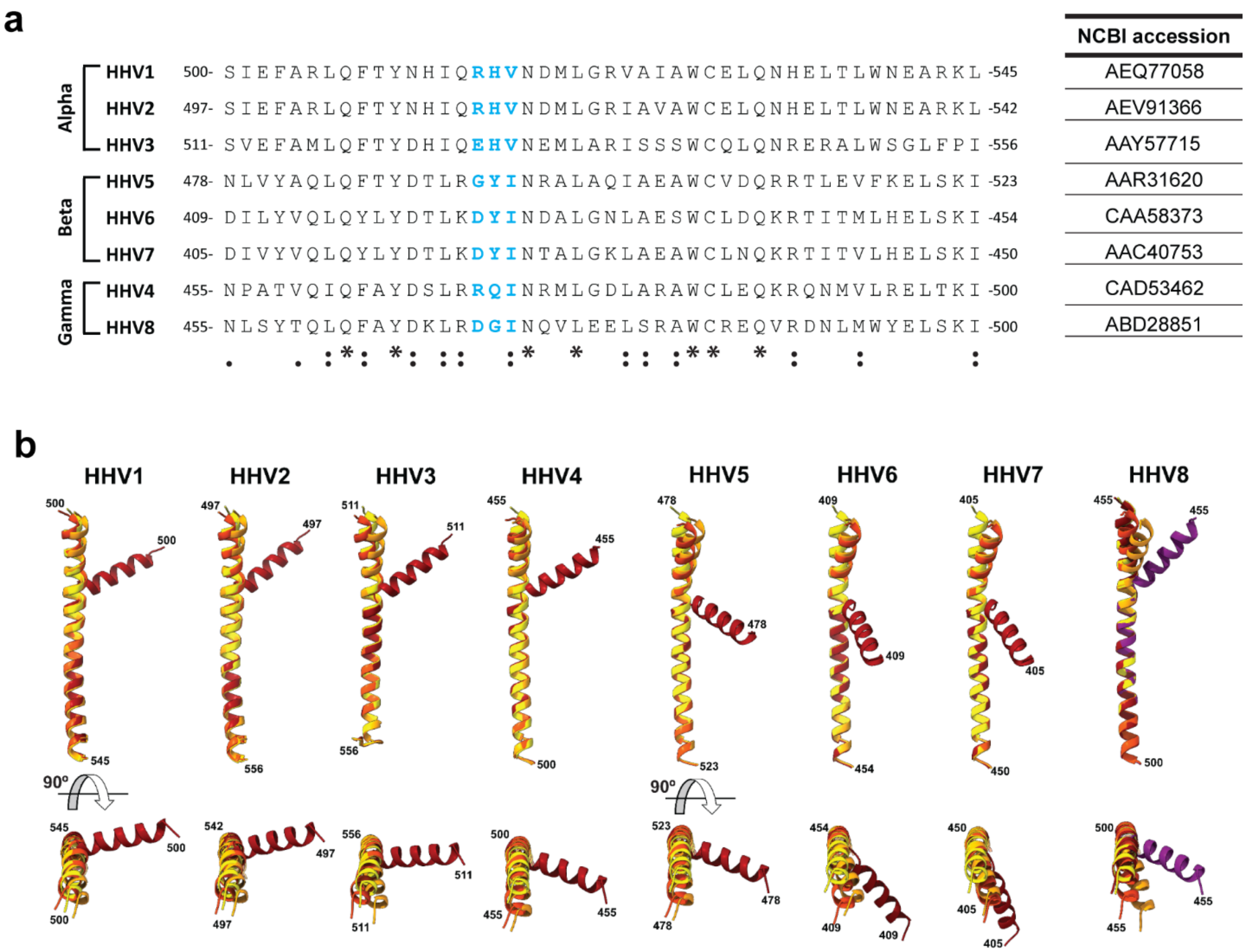

**Figure S6. Predicted structural deformations induced by proline substitutions to residues corresponding to VZV $^{526}$EHV$^{528}$ residues in gB DIII central helix of the eight human herpesviruses. a** Sequence alignment of the amino acids equivalent to VZV gB DIII central helix from human herpesviruses. gB NCBI accession are listed. Residues corresponding to VZV $^{526}$EHV$^{528}$ are colored in blue. Conservation among residues is presented based on Clustal Omega: "." indicates weak similarity; ":" indicates strong similarity; "*" indicates full conservation. **b** All structures for the wild type (WT) and mutant the gB DIII central helices were predicted using AlphaFold2 [3,4] and the mutant structures aligned to WT gB using Matchmaker [5]. WT gB is colored yellow and each of the proline substitutions is colored as for VZV gB E526 (orange), H527 (orange red), V528 (maroon). For HHV8, a fourth proline mutant, gB[G471Y/I472P] (purple) was modeled to assess the effect of tyrosine at the -1 position for I472, which had an RMSD of 7.109. The amino acids at the beginning and end of the gB DIII central helix are indicated. HHV1: HSV-1; HHV2: HSV-2; HHV3: VZV; HHV4: EBV; HHV5: HCMV; HHV8: KSHV.

**Table S1. Cryo-ET data collection parameters for WT gB and gB[H527P] EVs.**

| **Parameter** | **Structure** | | |
|---|---|---|---|
| | **WT gB[A]** | **gB[H527P][A]** | **gB[H527P][B]** |
| **Data collection and processing** | | | |
| Microscope | Titan Krios (FEI; D3690) | Titan Krios (FEI; D3690) | Titan Krios (FEI; D3690) |
| Voltage (kV) | 300 | 300 | 300 |
| Magnification (1000X) | 81 | 81 | 81 |
| Spot size | 6 | 6 | 5 |
| Energy filter slit width (eV) | 20 | 20 | 20 |
| Detector | Gatan K3 | Gatan K3 | Gatan K3 |
| Defocus range (μm) | -2.0 | -2.0 | -2.0 |
| Pixel size (Å) | 1.095 | 1.095 | 1.095 |
| Tilt axis (°) | 85.6 | 85.6 | 85.6 |
| Tilt range (°) | ± 60 (bi-directional) | ± 60 (bi-directional) | ± 60 (bi-directional) |
| Tilt increment | 3° | 3° | 3° |
| Exposure time (s) | 0.2 / 0.25 | 0.2 | 0.3 |
| Frames | 4 | 4 | 6 |
| Electron exposure rate ($e^-/Å^2/s$) | 11.9 | 11.9 | 10.73 |
| Dose per frame ($e^-/Å^2$/frame) | 0.573 | 0.573 | 0.54 |
| Dose per tilt ($e^-/Å^2$) | 3.44 / 4.3 | 3.44 | 3.22 |
| Total electron exposure ($e^-/Å^2$) | 141 / 176.3 | 93.973 | 132 |
| Tomograms collected (no.) | 10 | 17 | 43 |
| **Tomogram Reconstruction** | | | |
| Symmetry imposed | | | 3 |
| Final particle images (no.) | N/A | N/A | 2,074 |
| Map resolution (Å) | N/A | N/A | 12-15Å |
| FSC threshold | N/A | N/A | 0.143 |

[A] EVs produced using WT gB and gB[H527P] pCAGGs expression constructs.

[B] EVs produced using gB[H527P]-His pCAGGs expression constructs.

**Table S2. Comparison of the structure deviation for proline and alanine mutants in the VZV gB DIII central helix predicted with AlphaFold2.**

| **Mutant** | **RMSD (Pruned)[A]** | | **RMSD (All)[A]** | |
|---|---|---|---|---|
| | **Å** | **AA** | **Å** | **AA** |
| WT-DIII[B] | 0.693 | 35 | 2.672 | 46 |
| E526A[B] | 0.926 | 43 | 1.932 | 46 |
| E526P[B] | 1.019 | 35 | 2.691 | 46 |
| H527A[B] | 0.848 | 43 | 1.909 | 46 |
| H527P[B] | 0.964 | 26 | 3.428 | 46 |
| V528A[B] | 0.937 | 43 | 1.944 | 46 |
| V528P[B] | 0.812 | 24 | 9.115 | 46 |
| E526A[C] | 0.553 | 46 | 0.553 | 46 |
| E526P[C] | 0.693 | 39 | 1.481 | 46 |
| H527A[C] | 0.730 | 45 | 0.813 | 46 |
| H527P[C] | 0.636 | 38 | 1.831 | 46 |
| V528A[C] | 0.651 | 46 | 0.651 | 46 |
| V528P[C] | 0.334 | 26 | 10.673 | 46 |

[A] RMSD calculated for comparisons between WT-DIII and mutant gB DIII helices using MatchMaker [5] (Chimera) Needleman-Wunsch alignment algorithm (BLOSUM-62 matrix). The RMSD values are given for the pruned sequences and the complete helix (All).

[B] Wild type gB DIII (AA511-556; WT-DIII) and DIII mutant helices predicted with AlphaFold2 compared to the structure of gB solved by cryo-EM (PDB).

[C] DIII mutant helices predicted with AlphaFold2 compared to wild type gB DIII predicted with AlphaFold2.

**Table S3. Comparison of the structure deviation for proline mutants in the gB DIII central helix for all human herpesviruses predicted with AlphaFold2.**

| Virus | Alias | Strain[A] | Residue[B] [RMSD[C]] | | |
|---|---|---|---|---|---|
| HHV1 | HSV-1 | 17 | R515 [0.760] | H516 [0.617] | V528 [11.016] |
| HHV2 | HSV-2 | HG52 | R512 [0.857] | H513 [0.749] | V514 [9.117] |
| HHV3 | VZV | Dumas | E526 [1.535] | H527 [1.090] | V528 [8.612] |
| HHV4 | EBV | B95A | R470 [2.413] | Q471 [2.278] | I472 [9.099] |
| HHV5 | HCMV | Merlin | G493 [2.355] | Y494 [2.870] | I495 [14.969] |
| HHV6A | - | Uganda | D424 [2.254] | Y425 [2.915] | I426 [16.107] |
| HHV6B | - | Z29 | D424 [2.674] | Y425 [2.815] | I426 [16.416] |
| HHV7 | - | RK | D420 [2.656] | Y421 [3.297] | I422 [16.160] |
| HHV8 | KSHV | GK18 | D470 [4.178] | G471 [2.420] | I472 [0.395][D] |

[A] The strain used was the type species for each human herpesvirus.

[B] The residue in the gB DIII central helix substituted with proline.

[C] RMSD calculated for comparisons between the WT-DIII for each human herpesvirus and mutant gB DIII helices using MatchMaker (Chimera) Needleman-Wunsch alignment algorithm (BLOSUM-62 matrix). The RMSD values are given for the 46 amino acids for the entire helix.

[D] A fourth mutant was generated, G471Y/I472P, to demonstrate the effect of the glycine residue on the lack of gross structural change for the KSHV gB DIII I472P mutant. The KSHV gB[G471Y/I472P] mutant was predicted to have a RMSD of 7.109.

**Table S4. Yield efficiency of the FC-TEM experiment.**

| BAC | BAC transfected cells | GFP+ cells recovered by FC sorting | Cells screened by TEM | Virus replicating cells identified by TEM |
|---|---|---|---|---|
| pOka-TKGFP | $8.7x10^6$ | 87885 | 1500 | 1 |
| pOka-TKGFP-gB[E526P] | $8.7x10^6$ | 40000 | 500 | 1 |
| pOka-TKGFP-gB[H527P] | $8.7x10^6$ | 67675 | 2096 | 2 |
| pOka-TKGFP-gB[V528P] | $8.7x10^6$ | 46662 | 1026 | 1 |

**Movie S1. Cryo-EM maps of the two conformations of gB[H527P].** This movie shows the fitting of the homology modelled prefusion VZV gB into 9.0Å cryo-EM structure of HSV-1 gB[H516P] [6], followed by STA maps of class I and class II gB[H527P] molecule with each model showing MDFF of the homology model of the VZV gB [2] based on HSV-1 gB[H516P] [6] into the cryo-EM maps. The VZV gB domains are colored in DI (cyan), DII (green), DIII (yellow), DIV (orange), DV (red) and linker regions (hot pink).
